# A practive faeces collection protocol for multidisciplinary research in wildlife science

**DOI:** 10.1101/537803

**Authors:** Suvankar Biswas, Supriya Bhatt, Shrutarshi Paul, Shrushti Modi, Tista Ghosh, Bilal Habib, Parag Nigam, Gautam Talukdar, Bivash Pandav, Samrat Mondol

## Abstract

Faecal samples have become important non-invasive source of information in wildlife biology and ecological research. Despite regular use of faeces, there is no universal protocol available for faeces collection and storage to answer various questions in wildlife biology. We collected 1408 faeces from ten different species using a dry sampling approach, and achieved 94.87% and 86.02% success rate in mitochondrial and nuclear marker amplifications. We also suggest a universal framework to use the same samples for different use. This protocol provides an easy, quick and cheap option to collect non-invasive samples from species living at different environmental conditions to answer multidisciplinary questions in wildlife biology.

## Introduction

Non-invasive samples, in particular faeces have become a regular choice in wildlife biology, population monitoring and ecological research globally. Advantages of faecal sample-based wildlife research include easy sample collection, access to large sample size and spatio-temporal coverage. Historically, large scale use of faeces in wildlife biology started with dietary analysis of animals ^1^ but the introduction of advanced molecular tools added a new dimension to non-invasive research. These molecular tools have allowed biologists to investigate questions regarding population genetics ^2,3^, species distribution ^4^, demography ^5,6^, evolutionary biology ^7^ and wildlife forensics ^8^. In more recent time, faecal samples have been used in addressing various questions related to wildlife physiology including endocrinology and reproductive capacity ^9,10^, along with parasitology ^11,12^, disease dynamics ^13^ and conservation genomics ^14^. The sampling and storage demands of various questions in non-invasive wildlife research have led to a gradual development of faecal sampling and storage protocols. A number of logistical factors including collector’s safety, storage in the field, shipping samples from remote field areas with different environmental conditions etc. have been considered while gradual development of these protocols.

Over the years, a number of faeces collection and storage approaches has been used in wildlife research that are broadly classified into three categories: a) dry sampling (for example simple drying ^15^, silica preservation ^16^); b) wet sampling (ethanol collection ^17^; TNE and DMSO buffer ^18^; DETs solution ^19^; RNA later ^20^) and c) two-step approach ^21^ (see Table 1 for details). While all of these approaches have been used in wildlife research, they have several logistical limitations making their implementation in the field challenging. For example, sampling with silica beads has advantages in post-collection sample transport and storage ^22^ but is not cost effective as it requires large amount of silica beads to keep samples moisture free in humid areas. Similarly, ethanol preservation, the most widely used wet sampling approach is also expensive, require specific training to collect samples and is often problematic during sample shipping from remote areas ^21^. Currently no universal sampling protocol is available and very limited work has focused on testing faeces sampling and storage protocols to answer different questions in non-invasive wildlife research ^17,22,23^. Most of such experimentations were conducted on captive animals ^24^ or the studies were performed under favorable environmental conditions ^25^.

**Table 1.**
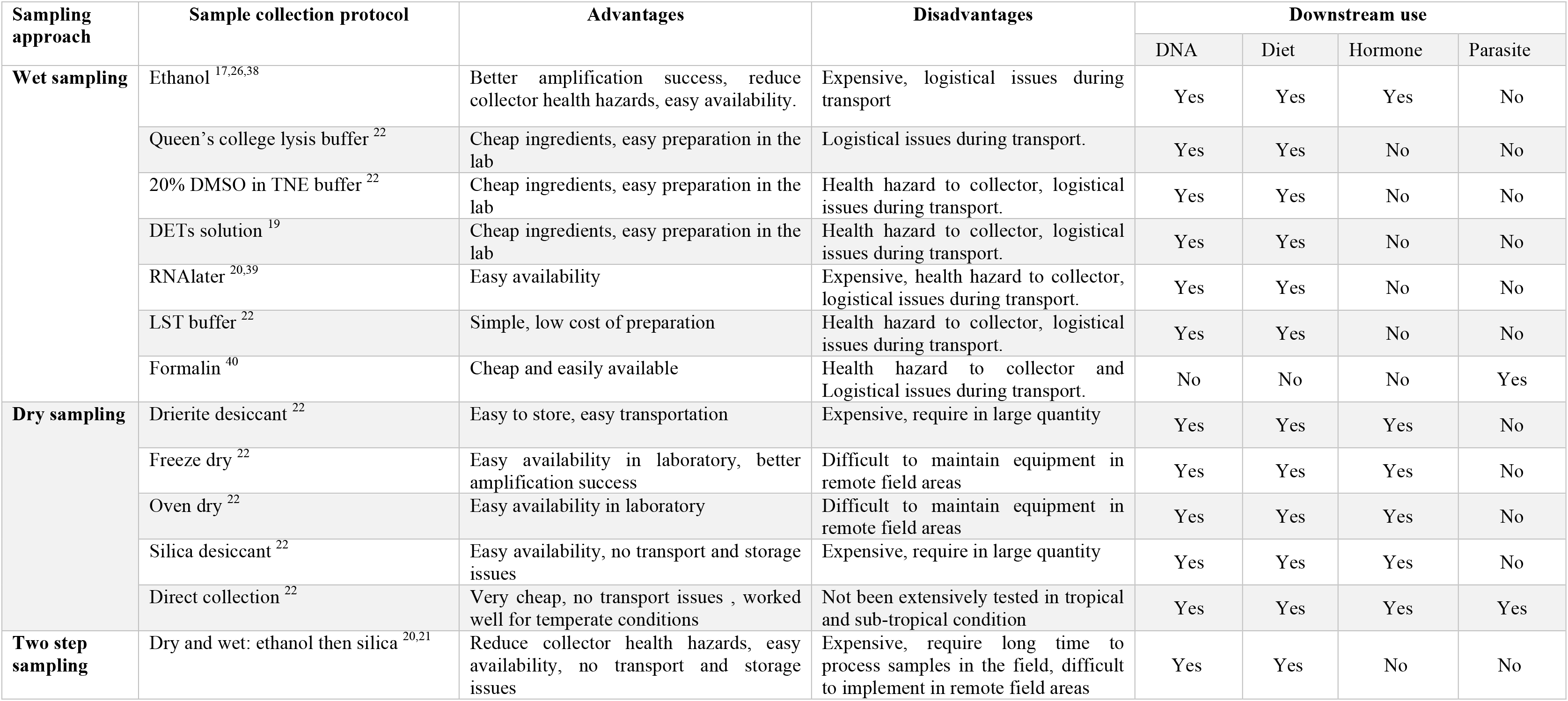
Details of different faecal sampling protocols and their downstream research use

In this paper, we describe a simple and cost effective dry sampling approach for faeces collection and storage that overcome the above-mentioned limitations and help answering different questions in faecal sample-based wildlife research. We followed this sampling approach to collect faecal samples of a number of carnivore and herbivore species living at different environmental conditions. Following sampling, we conducted molecular species identification and microsatellite amplification to demonstrate the efficacy of this approach for genetic work. Finally, we also propose a universal framework to use the faecal samples for various research purposes. We believe that the simplicity of the approach, ease of sample collection in the field and downstream use of the samples to answer various ecological questions will make this protocol useful in studying cryptic, elusive wildlife species across different environmental conditions globally.

## Methods

### Research permissions

All required permissions for our surveys and biological sample collections were provided by the Forest Departments of Uttarakhand (Permit no: 90/5-6 and 978/6-32/56), Uttar Pradesh (Permit no: 1127/23-2-12(G) and 2233/23-2-12 (G)) and Maharashtra (Permit no: 09/2016).

### Study habitats and species

In this study, our focus was to develop a faecal sampling protocol that could be used to answer different ecological questions (DNA, diet, parasite, hormone etc.) for terrestrial species. To test our protocol, we have collected samples from both herbivores (elephant, swamp deer, chital, Himalayan tahr) and carnivores (tiger, leopard, dhole, red fox, jungle cat, leopard cat), occupying various habitats ranging from sub alpine forest of lesser Himalayas, dry alpine scrub forests of trans Himalayan landscapes, moist-deciduous forests and swampy grass land of Terai-Arc landscape in north-western India and dry-deciduous forests of central Indian landscape. Sampling was conducted during different seasons across the states of Uttarakhand, Uttar Pradesh and Maharashtra where environmental conditions (ambient temperature, precipitation, humidity etc.) are varied.

### Collection and storage of faecal samples

We adopted a simple, cheap but effective field sampling protocol that involves cheap and easily available materials. Instead of standard use of absolute ethanol, silica gel, RNAlater or other similar approaches we collected faecal samples in butter paper (wax paper) and stored them individually into sterile zip-lock bags. The samples were stored inside dry, dark boxes in the field till they were transferred to the laboratory (within a maximum time of two months duration in this study). In the laboratory the samples were stored in −20° C freezers till further processing. All samples were collected with respective GPS locations and other associated field information.

We collected a total of 1408 faecal samples from 10 species across different habitats between December 2015 and May 2017. During collection the samples were categorised into respective species based on the morphological characteristics in the field. The details of species-wise sample sizes are provided in Table 2.

**Table 2.** Details of faecal samples collected and molecular methods used to test the efficacy of the dry sampling approach

| SI No. | Species name | Area/landscape | No. of samples<br>(Total N=1408) | DNA extraction protocol | Species confirmation method | Microsatellite loci used |
| --- | --- | --- | --- | --- | --- | --- |
| 1 | Tiger | Terai-Arc landscape, India | 1125 | Swab | PCR-Electrophoresis<sup>41</sup> | 13<sup>42 43</sup> |
| 2 | Leopard | Terai-Arc landscape, India |  | Swab | PCR-Electrophoresis<sup>41,43,44</sup> | 12<sup>42,43</sup> |
| 3 | Dhole | Central Indian landscape | 130 | Swab and scrape | PCR-Electrophoresis<sup>45</sup> | 04<sup>46 47</sup> |
| 4 | Red fox | Trans Himalayas | 05 | Swab | PCR-Sequencing<sup>48</sup> | 04<sup>46 47</sup> |
| 5 | Jungle cat | Lesser Himalayas | 15 | Swab and scrape | PCR-Sequencing<sup>48</sup> | 04<sup>42 43</sup> |
| 6 | Leopard cat | Lesser Himalayas | 06 | Swab and scrape | PCR-Sequencing<sup>48</sup> | 04<sup>42 43</sup> |
| 7 | Elephant | Terai-Arc landscape, India | 11 | Swab | NA | 03<sup>49</sup> |
| 8 | Swamp deer | Terai-Arc landscape, India | 85 | Swab | PCR-Sequencing<sup>50</sup> | 03<sup>51</sup> |
| 9 | Chital | Terai-Arc landscape, India | 22 | Swab | PCR-Sequencing<sup>50</sup> | 03<sup>51</sup> |
| 10 | Himalayan tahr | Middle Himalayas | 09 | Swab | PCR-Sequencing<sup>50</sup> | 03<sup>51</sup> |

### Faecal DNA extraction

To check the DNA quality following this dry sampling approach, we tested two different DNA extraction protocols in the laboratory. Both methods were initially tested with few faecal samples collected from different habitat types before employed in large scale sample processing. Our first approach was a slightly modified version of faecal swabbing protocol described in Ball et al., (2007) ^25^. This approach is advantageous over others as it retains most of the host cells from top layer and reduces the inhibitors present inside the faecal samples. Frozen faecal samples were thawed at room temperature and the upper layer was swabbed with Phosphate buffer saline (PBS) saturated sterile cotton applicators (HiMedia). Each sample was swabbed twice separately and were immediately stored in separate 2 ml microcentrifuge tubes in −20° C freezers till further processing. During extraction, 30 µl of Proteinase K (20mg/ml) and 300 µl of ATL buffer (Qiagen Inc.) were added into each tube containing swab and incubated overnight at 56° C, followed by Qiagen DNAeasy tissue DNA kit extraction protocol. DNA was eluted twice in 100 µl preheated 1X TE buffer. For every set of 22 samples two extraction negatives were taken to monitor any possible contaminations.

In the second approach, we scraped the top layer of faecal samples with sterile blade and stored in 2 ml microcentrifuge tubes for further processing. DNA extractions were performed using QIAamp DNA stool mini kit (QIAGEN Inc.) using protocol described in Mondol et al., (2009) ^26^. All faecal DNA extractions were conducted in a physically separated low-quality DNA extraction room.

### Molecular data generation

Field-collected non-invasive samples often generate low quantity and quality DNA for downstream molecular work ^27^. In this study, we tested efficacy of the sampling and DNA extraction protocols through molecular species identification (using mitochondrial DNA) and amplification of nuclear DNA (microsatellites) from faecal DNA samples collected in the field during this study.

#### 1) Species identification (using mitochondrial DNA)

We have adopted a number of approaches currently available for species identification from faecal samples of different species. These methods involved both species-specific PCRs as well as sequencing-based methods. The details of species specific approaches used for species identification are provided in Table 2. We did not perform molecular species identification for elephants due to morphologically distinctive appearances of its dung in field.

#### 2) Nuclear DNA (microsatellite) amplification

Amplification of nuclear DNA from non-invasive samples is challenging due to poor quantity and quality of DNA ^27^. In this study we have also amplified nuclear microsatellite markers from our field-collected faecal samples. We used a number of microsatellite markers to test the quality of DNA from field-collected samples from different species (see Table 2 for details). Species-wise cumulative amplification success rates for all tested loci were calculated.

## Results

We considered species identification and nuclear microsatellite amplification success rates from both swabbing and scraping protocols as efficacy of our faecal sampling approach for non-invasive wildlife genetic research. Initially we tested both approaches with 100 field-collected carnivore faecal samples (50 were swabbed and 50 were scraped) and achieved 100% success rates in species identification. As both approaches produced high success rates from field-collected faeces we compared other factors such as consumable cost, easeness of extraction protocol, time required etc. across both methods, and finally adopted the swabbing approach for the larger sample size. Subsequently, we swabbed the remaining 1308 faecal samples of different carnivore and herbivore species (see Table 2) collected from different habitats across India. Our overall success rate in species identification from all field-collected faecal samples (n= 1408) was 94.87%. Species-wise success rate details are given in Table 3.

**Table 3.** Species identification and microsatellite marker amplification success rates

| S. No. | Suspected species | No. of samples | Species identified samples | Success rate (%) | Samples with microsatellite loci data | Average success rate (%) |
| --- | --- | --- | --- | --- | --- | --- |
| 1 | Tiger | 1125 | 826 | 73.42 | 408 | 66.23 |
| 2 | Leopard |  |  |  | 159 | 62.5 |
| 3 | Dhole | 130 | 126 | 96.92 | 126 | 62.5 |
| 4 | Red fox | 05 | 05 | 100 | 05 | 69 |
| 5 | Jungle cat | 15 | 15 | 100 | 04 | 100 |
| 6 | Leopard cat | 06 | 06 | 100 | 04 | 100 |
| 7 | Elephant | 11 | 11 (NA) | 100 | 11 | 100 |
| 8 | Swamp deer | 85 | 71 | 83.53 | 71 | 100 |
| 9 | Chital | 22 | 22 | 100 | 11 | 100 |
| 10 | Himalayan tahr | 09 | 09 | 100 | 09 | 100 |
|  | Total | 1408 | 1091 | 94.87 | 821 | 86.02 |

Following species identification from field-collected faeces we targeted amplification of multiple nuclear microsatellites for different species. A total of 821 of 1091 samples from different species were successfully amplified for nuclear microsatellites, with a mean success rate of 86.02% (see Table 3 for details).

## Discussion

In this paper, we described a simple, fast and cost-effective faecal sampling approach for non-invasive wildlife research and tested this method on 10 different species that are found in a variety of different habitats. Development of field-suitable sampling and storage protocol is a progressive research in noninvasive wildlife research as faeces degradation in the wild is accelerated by exposure to various environmental conditions including sunlight (UV), humidity, temperature and rain and make it challenging to generate useful information for any target species. In comparison to other studies involving field-sampling and storage protocol standardization ^17,22–25^ we collected very large number of samples (n= 1408) from multiple species covering wide variety of habitats to test this protocol. Depending on the regions, the samples were stored in field conditions for up to two months before processing in the laboratory. High amplification success in species identification and nuclear marker amplification from field-collected samples indicate the efficacy of the approach for DNA-based research. To the best of our knowledge, this is the first study to use such large sample size from varied species to test faecal sampling and storage protocol. Given our sampling from wide variety of habitats and range of species with different ecology, we believe that this protocol would work well in other species living in different habitats across the globe. This approach is much cheaper than other available protocols (for example, silica gel, ethanol, RNAlater etc.), takes less time in field and doesn’t require specific training of field staff while implementation. However, we strongly suggest appropriate safety protocols (mask, gloves, protective gears etc.) during sample collection and processing for dry sampling approaches as exposure to potential pathogens is possible from dry faeces.

Testing two different DNA extraction protocols during this study provided very important insights on generating good quality DNA data from samples of different qualities. We performed swabbing and scraping DNA extraction approaches on a set of 100 carnivore samples. Carnivore scat samples were specifically chosen for standardization as they are difficult to generate data due to presence of prey ^28^. Given similar success rates achieved from both approaches and considering low consumable cost, extraction time and easeness swabbing was used for the remaining samples. While earlier studies have shown great efficiency of this approach ^24,25,29–31^, swabbing was mostly conducted with fresh (≤ 24 h), captive and frozen (≤ 0°C) faecal samples. Due to higher success rates with large number of faeces from multiple species in this study, we recommend the use of swabbing approach in future non-invasive genetic research. However, it is also important to point out that the scraping approach would be useful for comparatively older (≥ 2 weeks) faeces where outer layer is disturbed and for samples collected from very dry/dusty regions where swabbing the top layer is not feasible. Though we have tested both approaches with reasonably large number of carnivore faecal samples, any new study should test out both approaches with a few field-collected samples of the target species or decide on specific approach based on the sample conditions and physical characteristics (strata, dryness, availability of faeces top layer etc.) of the study area.

Another major advantage of this dry sampling approach is the ability to use the same samples to generate additional information apart from DNA data at species/individual levels. We propose a useful framework to showcase different use of the same samples in addressing various important biological questions in wildlife biology (see Figure 1 for details). For example, following swabbing/scraping for DNA, the frozen sample can be lyophilized to separate faecal powder and remaining prey hairs/plant products ^16^. Morphological analyses of these hairs/plant materials can provide information on dietary preferences ^32,33^. Similarly, the faecal powder could subsequently be used in understanding physiological parameters (stress ^34,35^, reproductive fitness ^9,10,36^, social dominance ^37^ and food preferences ^16^. During field sampling, a part of the faeces can be collected in formalin to study parasite abundance ^11^. In conclusion, our dry faecal sampling method provide an easy, cheap option to collect non-invasive samples from terrestrial wild animals. This universal protocol can be used to collect samples from species living at different environmental conditions and answer various questions related to genetics, genomics, physiology, diet, health etc. Along with other ecological information, these parameters would help developing informed conservation plans for any target species.

**Figure 1.**
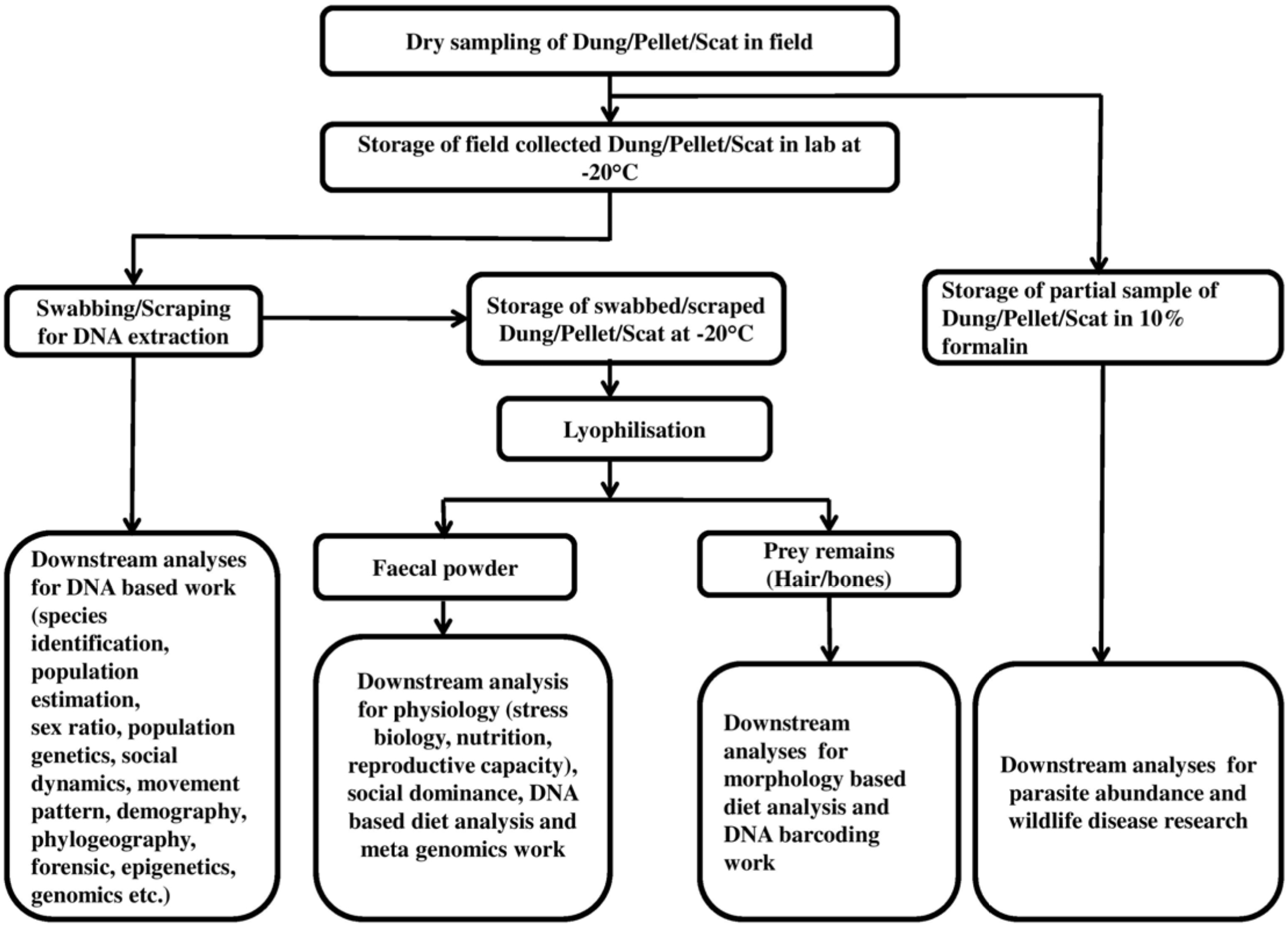
Flowchart showing a framework of various use of faecal samples collected through the dry sampling approach described in the manuscript.

## Acknowledgement

We acknowledge the Director, Dean and Nodal Officer of Wildlife Forensic Conservation Genetics Cell of Wildlife Institute of India for their support in this work. Our sincere thanks to Forest Departments of Uttarakhand, Uttar Pradesh and Maharashtra for research permits. Mr. A. Madhanraj and Ms. Zenab has provided critical support in the laboratory. We thank Dr. S. K. Gupta and Dr. S. P. Goyal for logistic support; Mr. H. S. Rathod and our field assistants Annu, Bura, Abbhi, Ranjhu and Imam for their effort in the field. This research was funded by Wildlife Conservation Trust-Panthera Global Cat Alliance Grants and Department of Science and Technology, Government of India. The corresponding author was supported by the INSPIRE faculty Award by the Department of Science and Technology.

